# Increased Pyramidal and VIP Neuronal Excitability in Primary Auditory Cortex Directly Correlates with Tinnitus Behavior

**DOI:** 10.1101/2022.11.22.517379

**Authors:** Madan Ghimire, Rui Cai, Lynne Ling, Kevin A. Brownell, Troy A. Hackett, Daniel A. Llano, Donald M. Caspary

## Abstract

Tinnitus affects roughly 15-20% of the population while severely impacting 10% of those afflicted. Tinnitus pathology is multifactorial, generally initiated by damage to the auditory periphery, resulting in a cascade of maladaptive plastic changes at multiple levels of the central auditory neuraxis as well as limbic and non-auditory cortical centers. Using a well-established condition-suppression model of tinnitus, we measured tinnitus-related changes in the microcircuits of excitatory/inhibitory neurons onto layer 5 pyramidal neurons (PNs), as well as changes in the excitability of vasoactive intestinal peptide (VIP) neurons in primary auditory cortex (A1). Patch-clamp recordings from PNs in A1 slices showed tinnitus-related increases in spontaneous excitatory postsynaptic currents (sEPSCs) and decreases in spontaneous inhibitory postsynaptic currents (sIPSCs). Both measures were directly correlated to the rat’s behavioral evidence of tinnitus. Tinnitus-related changes in PN excitability were independent of changes in A1 excitatory or inhibitory cell numbers. VIP neurons, part of an A1 local circuit that can disinhibit layer 5 PNs, showed significant tinnitus-related increases in excitability that directly correlated with the rat’s behavioral tinnitus score. That PN and VIP changes directly correlated to tinnitus behavior, suggests an essential role in A1 tinnitus pathology. Tinnitus-related A1 changes were similar to findings in studies of neuropathic pain in somatosensory cortex suggesting a common pathology of these troublesome perceptual impairments. Improved understanding between excitatory, inhibitory and disinhibitory sensory cortical circuits can serve as a model for testing therapeutic approaches to the treatment of tinnitus and chronic pain.

**Key points:**

- Identify tinnitus-related changes in synaptic function of specific neuronal subtypes in a reliable animal model of tinnitus.
- Finding show direct and indirect tinnitus-related losses of normal inhibitory function at A1 layer 5 pyramidal cells, and increased VIP excitability.
- Findings are similar to what has been shown for neuropathic pain suggesting that restoring normal inhibitory function at synaptic inputs onto A1 pyramidal neurons could conceptually reduce tinnitus discomfort.

## Introduction

Tinnitus, commonly referred to as ringing in the ears, is believed, in part, to reflect maladaptive plastic changes at various levels of the central auditory pathway as well as in non-auditory cortical areas, limbic and attentional structures. Partial deafferentation due to peripheral damage results in sensory deprivation to central auditory structures (Kujawa & Liberman, 2009; Shore & Wu, 2019; McGill *et al*., 2022). Canonical physiologic signs of decreased auditory input, described for tinnitus models, include increased spontaneous activity, increased neural synchrony and increased bursting at multiple levels of the central auditory pathway (Brozoski *et al*., 2002; Kaltenbach *et al*., 2004; Brozoski & Bauer, 2005; Kaltenbach *et al*., 2005; Ma *et al*., 2006; Schaette & Kempter, 2006; Bauer *et al*., 2008; Roberts *et al*., 2010; Noreña, 2011; Auerbach *et al*., 2014; Geven *et al*., 2014; Kalappa *et al*., 2014; Ropp *et al*., 2014). Markers for tinnitus-related maladaptive changes in neurotransmission are generally associated with disrupted excitatory glutamatergic and inhibitory GABA-/glycinergic signaling (Wang *et al*., 2009; Roberts *et al*., 2010; Kaltenbach, 2011; Richardson *et al*., 2012; Eggermont, 2013; Zhang, 2013; Sametsky *et al*., 2015; Caspary & Llano, 2017). The present study posited that primary auditory cortex (A1) layer (L) 5 pyramidal neurons (PNs) and vasoactive intestinal peptide (VIP) inhibitory neurons would show selective tinnitus-related changes in glutamatergic and GABAergic signaling.

Similar to somatosensory and visual cortices, the auditory cortex (A1) houses a complex local-circuitry of excitatory and inhibitory neurons, responsible for integration and higher order processing of sensory information (Winer & Lee, 2007; Gonchar *et al*., 2008; Letzkus *et al*., 2011; Schneider *et al*., 2014; Mesik *et al*., 2015). Deeper A1 layers are engaged in top-down modulation of ascending auditory information and integration of cortico-cortical information (Nelson *et al*., 2013; Williamson & Polley, 2019; Blackwell *et al*., 2020). In layer 5 of A1, excitatory pyramidal neurons (PNs) are found to interact in local circuits with at least 3 distinct types of inhibitory interneurons, parvalbumin (PV), somatostatin (SOM), and vasoactive intestinal peptide (VIP) neurons, located across layers (Blackwell & Geffen, 2017; Askew *et al*., 2019). Layer 5B PNs project excitatory glutamatergic information to subcortical nuclei, primarily inferior colliculus (IC), extra-lemniscal medial geniculate body (MGB) as well as to secondary auditory and non-auditory cortical structures (Barbour & Callaway, 2008; Stebbings *et al*., 2014; Sottile *et al*., 2017; Williamson & Polley, 2019). Apical dendrites of layer 5B PNs receive mono- and polysynaptic glutamatergic excitatory ascending signals from MGB and modulating inhibitory inputs from surrounding GABAergic interneurons (Letzkus *et al*., 2011; Schneider *et al*., 2014; Mesik *et al*., 2015; Blackwell & Geffen, 2017; Phillips *et al*., 2017; Askew *et al*., 2019; Blackwell *et al*., 2020). Within A1 excitatory inputs originate from the columnar principal neurons (Barbour & Callaway, 2008), VIP neurons regulate the excitability of layer 5 PNs through disinhibition (Pfeffer *et al*., 2013; Pi *et al*., 2013). Activation of PV and SOM interneurons provides direct inhibition onto PNs, while VIP GABAergic interneurons contact PV and SOM neurons, effectively disinhibiting PNs when activated (Letzkus *et al*., 2011; Pfeffer *et al*., 2013; Pi *et al*., 2013; Askew *et al*., 2019). The present study examined the impact of tinnitus on layer 5B PNs and VIP neurons. We used an established operant rat model of tinnitus in a series of whole-cell patch-clamp studies to investigate tinnitus-related synaptic changes in A1 layer 5B PNs and VIP neurons.

## Methods

### Animals

49 wild-type (WT, Envigo, Indianapolis, IN), 8 ChAT-Cre (Rat Resource & Research Center, Columbia, MO) and 19 *VIP^Cre^:Rosa26^tdTomato^* (Envigo, Indianapolis, IN) adult Long-Evans (LE) rats were entered into the studies at 3-4 mos. and used in a series of experiments at 8-10 mos. Experimental details of animal usages are shown in table 1. All animals were single housed, with a restricted diet schedule starting at the time of noise-exposure until the assessment of tinnitus. A reverse light/dark cycle was maintained, in order to conduct behavioral studies during active periods of the animals. Animals were handled according to SIU-School of Medicine LAB Animal Care and Use Committee (LACUC) approved protocols.

**Table 1:** Animals used in the study.

| Animals | Wild-type |  |  | ChAT-Cre |  |  | VIP x tdTomato |  |  |
| --- | --- | --- | --- | --- | --- | --- | --- | --- | --- |
|  | Control | ET | ENT | Control | ET | ENT | Control | ET | ENT |
| ABR threshold shift at 16 kHz<br>(Mean dB ± SD) | 3.34 ±<br>9.29 | 18.33 ±<br>11.25 | 17.5 ±<br>4.62 | 5.75 ±<br>9.39 | 11.4 ±<br>13.53 | 13.33 ±<br>13.53 | 2.0 ±<br>4.42 | 22.5 ±<br>7.5 | 13.33 ±<br>11.24 |
| Electrophysiological studies<br>(Rats/Neurons) | 16/62 | 19/77 | 2/12 | 3/10 | 5/14 | 6/25 | 7/28 | 6/22 | 0 |
| Anatomical studies | 7 | 5 | 2 | 0 | 0 | 0 | 3 | 0 | 3 |
| Total animals used | 23 | 24 | 4 | 3 | 5 | 6 | 10 | 6 | 3 |

### Tinnitus Induction

To assure that all animals used in the present study had normal hearing thresholds, before noise-exposure, animals were anaesthetized using a cocktail of ketamine (90 mg/kg)/xylazine (7 mg/kg) and auditory brainstem response (ABR) thresholds were obtained before and after the exposure for pure tones at 8, 10, 12, 16, 20, 24, and 32 kHz, presented in 10 dB increments between 10 and 80 dB (SPL re 20 μPa). Unexposed animals were similarly tested (Fig. 1A). Briefly, 3-4 mo WT, ChAT-Cre or *VIP^Cre^:Rosa26^tdTomato^* LE rats were classified as entering the control or exposure groups. Two thirds of the animals were unilaterally noise-exposed as in prior studies by Bauer and Brozoski (Bauer *et al*., 1999; Bauer & Brozoski, 2001). The sound-exposed group was divided into animals with behavioral evidence of tinnitus (ET) or without significant behavioral evidence of tinnitus (ENT). Remaining unexposed animals were identically treated but were not noise-exposed and are referred to as controls. Noise-exposure was similar to previous studies conducted on rats (Bauer & Brozoski, 2001; Brozoski *et al*., 2013). One week following ABR testing, at 3 months of age, rats were anesthetized with a 1.7% isoflurane/O_2_ mixture and placed in a head holder for noise-exposure of the right ear. Rats were unilaterally exposed to narrow-band (16 kHz centered) noise for 1 hour, with a peak level of 116 dB (SPL), falling to ambient levels at 8 kHz and 24 kHz, with the contralateral canal blocked (Bauer & Brozoski, 2001, 2006). ***Calibration:*** Prior to each set of noise-exposures/ABRs, subsets of *in vivo* and behavioral experiments, sound levels were calibrated using a Brüel and Kjaer Pulse sound measurement system (13.1 software; 4138 microphone) (see Kalappa *et al*., 2014). ***Tinnitus Assessment:*** Three months following noise-exposure, tinnitus was measured, using a behavioral assay sensitive to tinnitus in rats, and detailed in Bauer and Brozoski (Bauer & Brozoski, 2001; Galazyuk & Brozoski, 2020) and Brozoski et al. (Brozoski *et al*., 2012). Briefly, an operant conditioned-suppression procedure determined the animal’s perception of test tones and silent periods embedded in an ambient, low level (60 dB, SPL) broad-band noise (BBN). Animals were required to discriminate between the presence and absence of sound and were tested daily in modified operant test chambers. Foot shock provided negative reinforcement to the animals and quickly shaped their behavior. Animals able to identify silence suppressed lever pressing during the silent period, however, animals unable to identify silence from tone period would suppress lever pressing during both silence and tone period. Based on their lever press, suppression ratio was calculated using the formula: R = A/(A+B) where R = Suppression ratio; A = lever press during 16/20 kHz tone period, and B = lever press following (two min bin) tone period. Three criteria had to be met for individual-subject data to be included for further analysis: 1) There had to be a minimum of 200 lever presses in each session; 2) mean *R* for background noise periods (i.e., baseline performance) had to be > 0.4, with 3) a SD of < 0.2. Data from a minimum of five test sessions, for each stimulus, were averaged to derive individual and group discrimination functions for each stimulus: BBN, 16, and 20 kHz tones. Control (Ctrl) animals showed higher suppression ratios while animals showing evidence of tinnitus had lower suppression ratios demonstrating their impaired ability to identify pure tone (Fig. 1B). The noise exposed animals were classified as exposed tinnitus (ET) if the average suppression ratio was lower than 0.26 and animals with average suppression ratio higher than 0.26 were classified as exposed non-tinnitus (ENT) at 16kHz discrimination task. Once psychophysical tests were completed, animals’ exit ABR were collected prior to *in vitro* studies.

**Figure 1:**
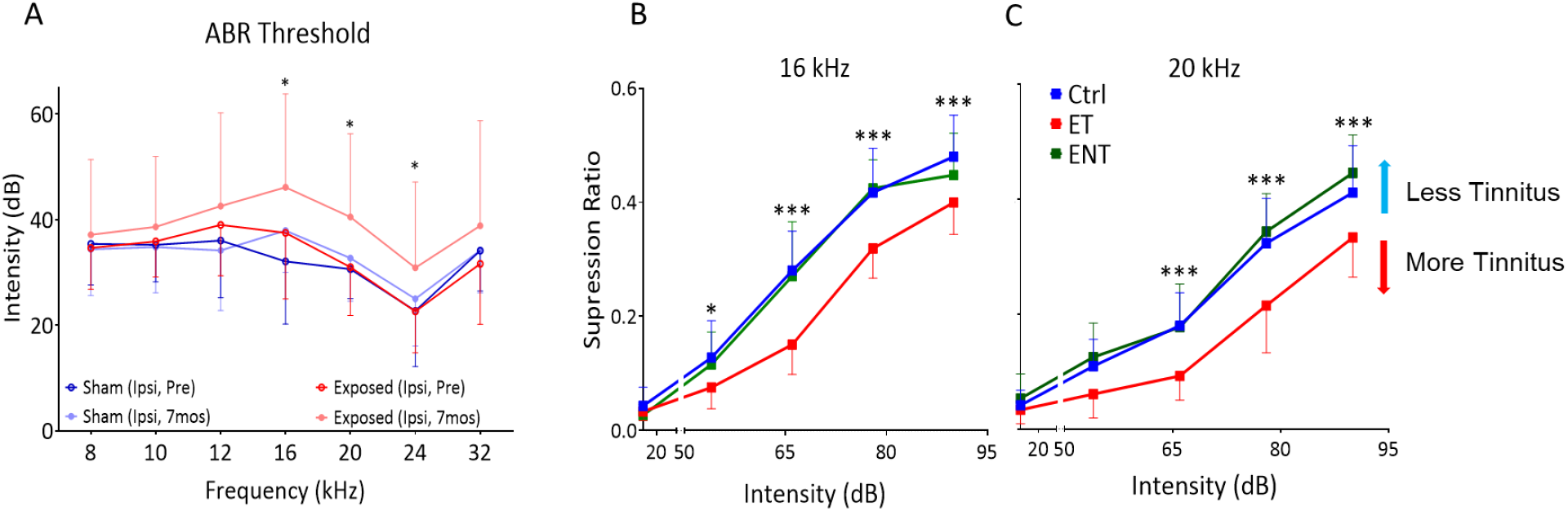
Animal model of tinnitus. A. ABR threshold before and after unilateral noise-exposure in sham exposed controls (n = 40) and noise-exposed animals (n = 24). A modest significant shift in ABR threshold after noise-exposure was observed (F (1, 554) = 25.28, *p* = 0.0001, Two-way ANOVA). B, C. Based on their suppression ratios, animals trained on the discrimination task were categorized as Control (Ctrl, n = 22), exposed tinnitus (ET, n = 15) and exposed non-tinnitus (ENT, n = 13). A significant separation between the suppression ratios of Ctrl and ET was observed at 16 kHz (B) (F (2,230) = 32.28, *p* < 0.0001, Bonferroni post-hoc test) and 20kHz (C) (F (2,230) = 35.77, *p* < 0.0001, Bonferroni post-hoc test) frequency tests. * *p* < 0.05, *** < 0.001.

### Micro-injection surgery

Tinnitus tested wild-type and *VIP^Cre^:Rosa26^tdTomato^* LE rats (8-9 mos) were anaesthetized with a cocktail of ketamine (90 mg/kg)/xylazine (7 mg/kg) (induction anesthesia) followed by 0.5-1% isoflurane (maintenance). Animals were head fixed in a stereotaxic apparatus with viral vectors injected intracranially using a Neurostar stereotaxic drill and injection system (stereodrive 015.838, injectomate IM28350, stereodrill DR352; Neurostar, Germany). A midline incision was made to expose the skull. In wild-type rats, a craniotomy of 1.5 mm was drilled at AP −5.7 and ML −3.5, corresponding to MGB injection site. A 2 μl Hamilton micro-syringe was used to inject approximately 250-300 nl of AAV5-CaMKIIa-hChR2(H134R)-EYFP (UNC Vector Core, UNC, Chapel Hill, NC) 6mm ventral to bregma. In *VIP^Cre^:Rosa26^tdTomato^* rats, a craniotomy of 1.5 mm was drilled at AP −5.7 and ML −4.9 respective to bregma corresponding to A1 injection site. A 2 μl Hamilton micro-syringe containing AAV-EF1a-DIO-hChR2(H134R)-EYFP (UNC Vector Core, UNC, Chapel Hill, NC) was fixed at −22-degree angle in a sagittal plane. The syringe was advanced 5 mm from the surface of the brain to reach A1, and approximately 150-200 nl of the virus was injected. Animals were allowed to recover and express the virus for a minimum of 3 weeks and were used in *in vitro* optogenetic studies.

### Cortical slice electrophysiology

LE rats (8-9 mos) were anesthetized with 4% isoflurane and cardiac perfusion was performed using ice-cold sucrose aCSF (in mM as follows: 2.5 KCl, 5 MgCl_2_, 1.23 NaH_2_PO_4_, 0.5 CaCl_2_, 250 sucrose, 26 NaHCO_3_, and 10 glucose, pH 7.4) saturated with carbogen (95% O_2_/5% CO_2_) before decapitation. After perfusion and decapitation, brains were rapidly isolated and submerged in ice-cold (1°C – 2°C) sucrose aCSF, pH ~7.4, and oxygen saturation was maintained by bubbling with carbogen. aCSF composition was as follows (in mM): 125 NaCl, 3 KCl, 1 MgCl_2_, 1.23 NaH_2_PO_4_, 2 CaCl_2_, 26 NaHCO_3_, and 10 glucose. Coronal slices of 250–300 μm through A1 were sectioned using a vibratome (Pelco), following previously described protocols (Richardson *et al*., 2013; Ghimire *et al*., 2020), and incubated for 15 min at 31°C. Slices were allowed to equilibrate at room temperature (20°C – 22°C) for 60 min in carbogen-bubbled aCSF before recordings. Slices were transferred to an immersion recording chamber (2 ml), perfused at 2–3 ml/min with aCSF bubbled with carbogen, and all the recordings were performed at room temperature. Layer 5 PNs were identified morphologically using QImaging Rolera bolted on a differential interference contrast microscope (BX50WI, Olympus Optical) under a 45× water-immersion objective. Likewise, tdTomato labelled VIP neurons were identified using fluorescence system (lambda 421, Sutter Instruments) with excitation 560 nm/emission 581 nm, attached to the DIC microscope.

### *In vitro* patch-clamp recordings

Once cells were identified, whole-cell patch-clamp recordings were performed using 3-6 mΩ fire-polished microfilament micropipettes, pulled from borosilicate glass (0.86 mm ID, 1.5 mm OD; Sutter Instruments). Two different composition of the internal solution were used 1) (in mM): 140 potassium gluconate, 1 NaCl, 2 MgCl_2_, 5 KCl, 10 HEPES, 2 Mg-ATP, 0.3 Na-GTP, 6.88 KOH, Osm: ^~^300 mOsms, pH 7.3 (adjusted with Tris-Base) with calculated chloride reversal potential near −65 mV and 2) (in mm) 140 CsCl, 2 MgCl2, 4 Mg-ATP, 0.3 Na-GTP, 10 Na-HEPES, 5 QX 314 and 0.1 EGTA, with a pH of 7.25 adjusted with HCl (osmolarity, ~290 mOsm), resulting in a Cl-equilibrium potential (ECl–) value near 0 mV. Pipettes were attached to multi-clamp 700B Amplifier (Molecular Devices), and cells recorded in current-clamp mode at I = 0 or voltage-clamp mode at −65 mV at 10 kHz sampling rate. The patch pipette was attached to layer 5 PNs or layer 2/3 VIP neurons with Giga-ohm (> 4 GΩ) seal and the membrane was ruptured with a brief sharp suction. Whole-cell recordings were collected with access resistances ranging from 10-25 mΩ. Whole-cell capacitance, input resistance, and access resistance were determined by injection of a 5-mV square pulse, at 20 Hz. Exclusion criteria included the following: 1) a resting membrane potential more depolarized than −55 mV, 2) access resistance > 25 mΩ, 3) a resting input resistance < 100 mΩ (for PNs only). TTL pulses, voltage commands, acquisition, and display of the recorded signals were achieved with Axon Digidata 1440A (Molecular Devices) using the Clampex 10.7 program.

### Optogenetic studies

Optogenetic studies used adult (8-10 mos) wild-type and *VIP^Cre^:Rosa26^tdTomato^* rats that underwent viral injection at MGB or A1. 10-20 ms pulses of blue light (460 nm) were used to activate ChR2 with a Lambda 421 optical beam combiner system (Sutter instruments) illuminating area of interest through microscope objective lens. The power of the LED was set to constant at 1 Amps, and optical stimulation was made at focus distance of the microscope at 45X objective.

### Saturation Binding

Binding protocols were identical to those used previously (Richardson *et al*., 2011; Ling & Caspary, 2013). Five control and five noise-exposed LE rats (8-10 mos) were decapitated, and the brains were rapidly isolated, rinsed in ice-cold phosphate buffer at 4°C, pH 7.4, frozen in powdered dry ice, and stored at −80°C. Serial transverse sections were cut at 16 μm using a Leica CM1850 cryostat at −24°C. Selected sections were thaw-mounted onto Superfrost/Plus slides and stored at −20°C. Anatomical locations of the A1 were verified to match neural structures with those described previously (Paxinos & Watson, 1998).

[^3^H]gaboxadol (a gift from Merck, Inc.) was used with modified protocols from Dr. Bjarke Ebert (H. Lundbeck A/S, Copenhagen, DK; personal communication) as described in Richardson et al. (2011). After fixed in 4% PFA for 10 minutes at room temperature, tissue sections were subjected to prewash twice for 5 min in buffer, followed by incubation with [^3^H] gaboxadol: 10–400 nM for 1 hour at 4°C and post-wash with buffer for four quick dips. The buffer solution used was 50 mM Tris-citrate, pH 7.1. Nonspecific binding was determined in adjacent sections by the addition of cold excessive GABA to the ligand buffer and was subtracted from total binding.

Dried slides were opposed to [^3^H]-hypersensitive phosphor screens for 2 d at room temperature. The phosphor screens were scanned using a Cyclone Storage Phosphor System (PerkinElmer). The layer 2-6 of A1 was outlined and analyzed using OptiQuant Image Analysis software (Canberra Packard), which provided tools for grayscale quantification in digital light units. Digital light units were then converted to nanocuries per milligram protein using a standard curve generated from co-exposed [3H]-embedded plastic standards (American Radiolabeled Chemicals). GraphPad Prism software was used for data analysis and generating graphs.

### Cellular phenotyping and quantification

Multiplexed fluorescence in situ hybridization (FISH) was used to identify neuronal cell types in fresh frozen tissue sections (14 μm) across AI layers. Cells were identified by detection of cell-type-specific markers for excitatory neurons (VGluT1, Slc17a7), and inhibitory neurons (VGAT, Slc32a1) with their subtypes (parvalbumin, Pvalb; vasoactive intestinal peptide, Vip; somatostatin, Som) in the same tissue sections, counterstained with DAPI. Assays were conducted in sections collected from primary auditory cortex (A1) of 5 animals in each experimental group using custom RNAscope riboprobes and detection kits manufactured by Advanced Cell Diagnostics (Newark, CA), as in prior studies from our lab (Ghimire *et al*., 2020). Sections were imaged at 20x using a Nikon 90i epifluorescence microscope, permitting selective detection of transcripts for each cell marker by color channel. Images were imported in HALO pathology software (Indica labs, Albuquerque, NM) for analysis. Cells expressing the transcripts of each cell marker were separately tallied in each layer of A1 (Table 2).

**Table 2:** Tinnitus-related changes in cell density across the layers of A1.

| Cell Type | L 2/3 |  | L4 |  | L5 |  | L6 |  |
| --- | --- | --- | --- | --- | --- | --- | --- | --- |
|  | Control | Tinnitus | Control | Tinnitus | Control | Tinnitus | Control | Tinnitus |
| VGLUT 1 (cells/mm<sup>2</sup>) ± SD | 1267.49 ± 284.37 | 1320.78 ± 200.45 | 1142.74 ± 207.09 | 1180.18 ± 122.60 | 781.46 ± 268.4 | 878.82 ± 154.7 | 953.54 ± 497.64 | 1060.34 ± 179.34 |
| VGAT (cells/mm<sup>2</sup>) ± SD | 262.10 ± 79.2 | 263.14 ± 69.77 | 380.35 ± 68.22 | 289.74 ± 65.81 | 257.27 ± 75.06 | 256.04 ± 43.98 | 197.50 ± 49.8 | 164.15 ± 32.57 |
| PV (cells/mm<sup>2</sup>) ± SD | 113.68 ± 43.98 | 116.72 ± 32.10 | 253.08 ± 89.42 | **174.59 ± 25.42\*** | 108.26 ± 16.36 | 126.69 ± 27.35 | 61.13 ± 24.37 | 42.58 ± 4.18 |
| VIP (cells/mm<sup>2</sup>) ± SD | 65.32 ± 17.65 | 67.52 ± 19.26 | 30.04 ± 14.69 | 28.8 ± 18.38 | 16.87 ± 6.03 | 15.03 ± 12.44 | 11.07 ± 5.17 | 7.29 ± 4.93 |
| SOM (cells/mm<sup>2</sup>) ± SD | 29.15 ± 10.7 | 38.68 ± 13.18 | 80.16 ± 36.56 | 79.35 ± 20.0 | 102.22 ± 43.10 | 86.97 ± 40.28 | 49.33 ± 16.21 | 59.04 ± 31.98 |

\* $p < 0.05$ , Two-way ANOVA with Bonferroni *post-hoc* correction.

### Statistical and data analysis

Signals obtained from electrophysiological recordings were filtered at 5 kHz lowpass Gaussian filter and sEPSCs were defined as spontaneous glutamatergic currents greater than 8 pA and sIPSCs were defined as spontaneous chloride currents greater than 10pA. Data were reduced using Clampfit 10.7 (Molecular Devices), followed by further sorting using a custom MATLAB program. Correlation across control and ET animals in different groups (entering studies at different times) was established using z-score as a function of tinnitus score. z-scores (z-score = (observed value-mean)/standard deviation) were calculated when tinnitus scores from tested animals in different entering groups were combined. Statistical analysis was performed using GraphPad Prism 9, with corresponding statistical tests matched to the data noted in the figure legends. Student’s *t*-tests were used for individual sample comparison and a one-way/two-way ANOVA with a Bonferroni *post hoc* correction was used for multiple comparisons. Data were tested for the homogeneity of variance, and *p* values corrected based on statistical significance. All comparisons with *p* value < 0.05 were considered significant.

## Results

Subjects in the present study were tested for changes in their acoustic threshold using the auditory brainstem-evoked response (ABR). ABR threshold changes in unexposed/control rats were compared with right ear/unilaterally sound-exposed Long Evans (LE) rats, which showed modest threshold shifts at frequencies adjacent to the 16 kHz exposure frequency at the end of the tinnitus test paradigm (3-4 months after exposure) (Fig. 1A). As detailed in the methods, noise-exposed animals were parsed into two groups (exposed tinnitus (ET), exposed non-tinnitus (ENT)) based on their behavioral suppression ratios indicative of psychophysical evidence of tinnitus (Fig. 1B, C). Tinnitus suppression scores were converted to z-scores to normalize suppression ratios across different sets/groups of rats studied over the course of the 3 years. Tinnitus testing occurred 3-4 months following noise-exposure (6-7 mos. of age) with 50-60% of noise-exposed rats showing behavioral evidence of tinnitus. Approximately 90% of control/unexposed animals showed near-base-level suppression ratios/non-tinnitus behavior (Fig. 1B, C). The third group of noise-exposed rats showed control-like behavior suggesting little evidence of tinnitus and were classified as exposed non-tinnitus (ENT) (Fig. 1B, C). *In vitro* recordings were obtained from whole-cell patch-clamped neurons (200 PNs and 50VIP interneurons) in A1 slices from 30 ET, 11 ENT and 26 unexposed-Ctrl rats (Table 1). A1 cell counts were obtained using fluorescence *in situ* hybridization from 5 control and 5 ET animals.

### A1 layer 5 PNs showed tinnitus-related increases in activity of glutamatergic input neurons

Whole-cell patch-clamp electrophysiology from 9 control, 11 ET, and 4 ENT rats was used to examine tinnitus-related changes in the physiology of A1 layer 5B PNs (Fig. 2A). The chloride reversal potential was −65 mV, voltage clamp recordings were collected at −65 mV with current clamp recordings collected at I = 0. Once whole-cell configuration was achieved, intrinsic properties including resting membrane potentials (RMP) and rheobase current (RC) were measured after the cell stabilized, typically after 2-3 minutes. Resting membrane potential of layer 5 PNs was found to be significantly depolarized in ET animals, while no measurable difference observed in ENT animals compared to control PNs (Fig. 2B). There were no tinnitus-related changes in rheobase current (RC, current required to evoke an action potential) between groups (Mean ± SD pA, Control, 74.33 ± 26.48, n = 30; ET 73.26± 29.98 n = 43; ENT, 87.14 ± 47.51, F(2, 77) = 0.0.63, *p* = 0.53, graph not shown). To assess the activity of excitatory input neurons, we collected continuous voltage-clamp recordings of the frequency and amplitude of depolarizing sEPSCs from layer 5 PNs (Fig. 2C). Layer 5 PNs from ET rats showed significant tinnitus-related increases in sEPSC frequency, with no significant sEPSC frequency changes observed in ENT rats (Fig. 2D). ET animals showed a significant linear relationship between the sEPSC frequency and normalized tinnitus score (z-score). This relationship was not significant for Ctrl and ENT animals (Fig. 2E). No significant changes in sEPSC peak amplitude was observed from the PNs of control, ET and ENT animals (Fig. 2F).

**Figure 2:**
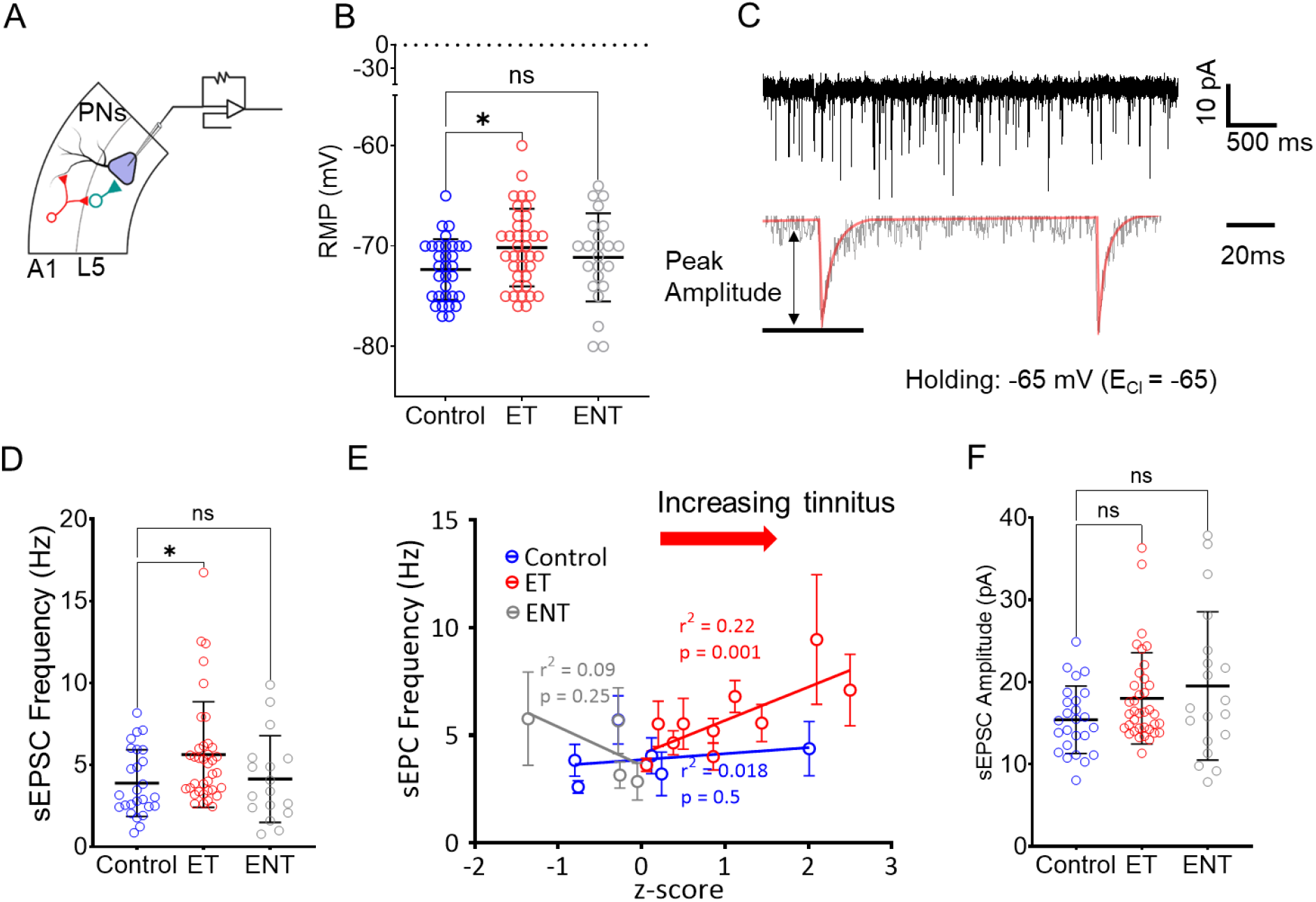
Tinnitus-related increases in activity of excitatory input neurons onto layer 5 PNs. A. Illustration representing patch-clamp recording configuration from A1 layer 5 PNs. B. Resting membrane potential (RMP) of PNs from ET animals was depolarized relative to layer 5 PNs from control neurons (Control, Mean ± SD mV, −72.34 ± 3.02, n = 32; ET, 70.16 ± 3.85 n = 38; ENT, 71.13 ±4.39, n = 23; F (2, 90) = 2.96, *p* = 0.0336, One-way ANOVA with Bonferroni post-hoc test). C. Top, exemplar recording trace of continuous voltage-clamped sEPSC recordings, bottom, measurement of peak amplitude. D. Tinnitus-related increase in frequency of sEPSCs recorded from layer 5 PNs from ET animals (Control, Mean ± SD Hz, 3.88 ± 2.03, n = 26; ET 5.62 ± 3.22 n = 37; ENT, 4.13 ± 2.64, n = 17; F (2, 77) = 3.56, *p* = 0.032, One-way ANOVA with Bonferroni post-hoc test). E. Simple linear regression analysis was performed to predict sEPSC frequency based on normalized tinnitus score(z-score^#^) (Control, F(1,24) = 0.44, r^2^ = 0.018, *p* = 0.25; ET, F(1,44) = 12.4, r^2^ = 0.22, *p* = 0.001; ENT, F(1, 14) = 1.4, r^2^ = 0.09, *p* = 0.25). F. Amplitude of sEPSCs recorded from layer 5 PNs in control ET and ENT animals (Control, Mean ± SD pA, 15.42 ± 4.1, n = 25; ET, 18.02 ± 5.5, n = 37; ENT, 19.52 ± 9.015, F (2, 78) = 2.56, *p* = 0.083, t-test). * *p* < 0.05. ^#^ See methods for z-score normalization.

### A1 layer 5 PNs show tinnitus-related increases in thalamocortical excitability

Layer 5 PNs receive glutamatergic inputs from thalamocortical neurons and indirect excitatory connections from within and outside A1 (Huang & Winer, 2000; Banks & Smith, 2011). MGB is the primary source of ascending excitatory auditory input to A1 terminating throughout A1 layers with largest projections to layers 1 and layer 4 (Huang & Winer, 2000; Banks & Smith, 2011; Takesian *et al*., 2018). Apical dendrites emerge from layer 5 PNs and ascend throughout granular and supragranular layers of A1 while receiving mono and polysynaptic thalamocortical endings (Slater *et al*., 2019). Tinnitus-related changes in the ascending MGB inputs onto layer 5 PNs were examined using optical stimulation of thalamocortical terminals in A1 (Fig. 3A). We injected AAV-ChR2 in the MGB and allowed the virus to express for 3-4 weeks. Acute slices of (~ 250-300 μm) were prepared and whole-cell patch-clamp recordings were performed. The expression of virus in the MGB neurons was confirmed by imaging the slices containing MGB and A1 for the presence of fluorophore EYFP (Fig. 3B). Thalamocortical endings appeared most prominent in layers 1 and 4 of A1 (Fig. 3B). Optical stimulation of thalamocortical terminals evoked significantly larger depolarizing currents (EPSCs) from PNs in ET animals with behavioral evidence of tinnitus than from PNs from control animals (Fig. 3C, D). TTX was then added to the bath in order to block polysynaptic thalamocortical responses. TTX significantly reduced the amplitude of thalamocortical-evoked EPSCs recorded from layer 5 PNs in both tinnitus and control animals (Fig. 3C, D). The present TTX findings are consistent with previous studies showing that much of the ascending excitatory drive from MGB reaches layer 5 PNs indirectly via local excitatory micro-circuitry (Slater *et al*., 2019). Observation of tinnitus-related increases in thalamocortical excitability prior to TTX blockade of polysynaptic thalamocortical responses and no changes in monosynaptic responses are suggestive of tinnitus-related plastic changes in both local A1 excitatory neurons and likely the projecting thalamocortical input.

**Figure 3:**
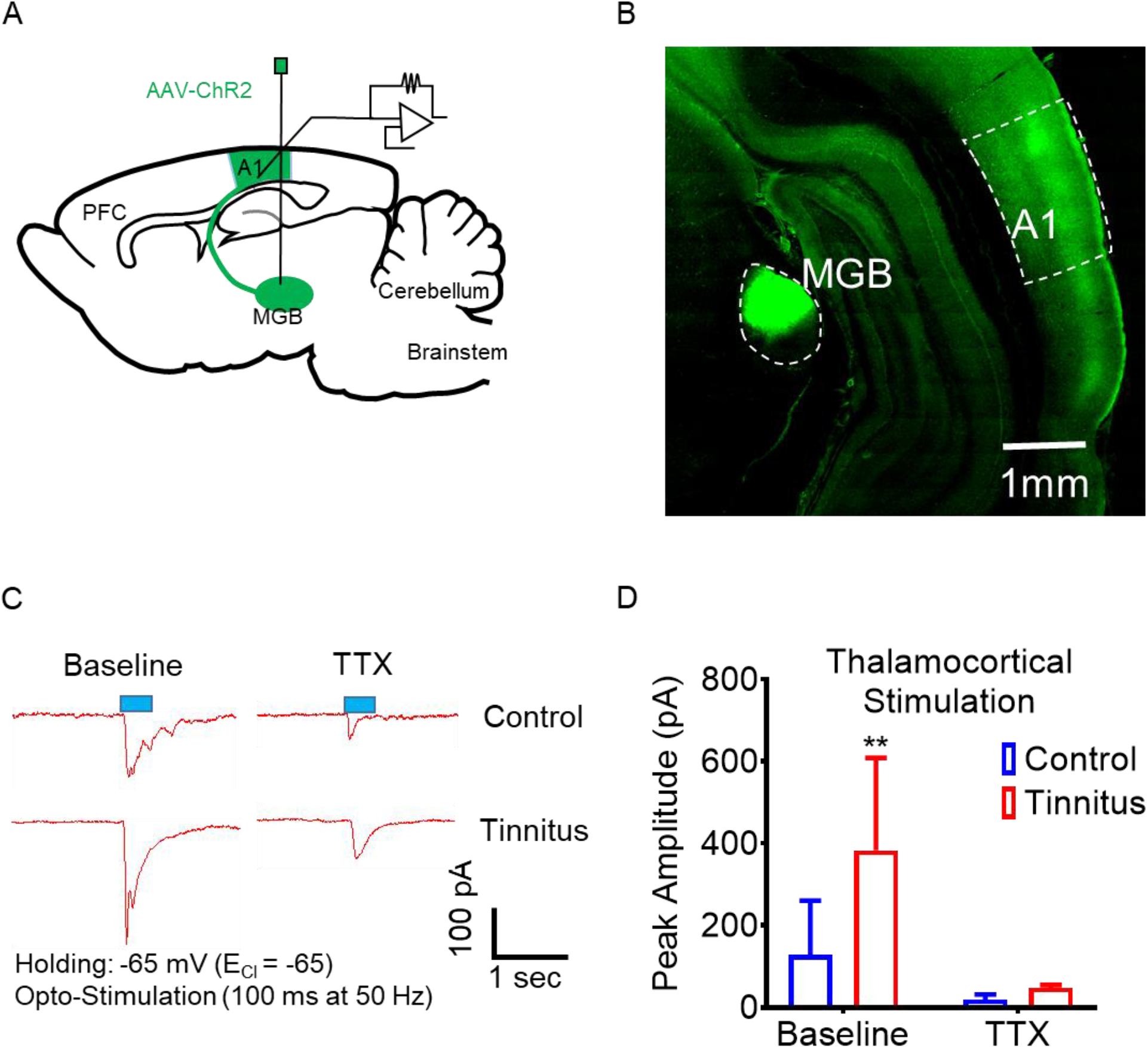
ET animals show increased thalamocortical excitability. A. Illustration showing injection of AAV-ChR2 in the MGB of control and ET animals. B. MGB principal projections into layer 1 and 4 of A1. C. Exemplar traces from control and ET animals showing evoked sEPSCs from layer 5 PNs evoked by optical stimulation of MGB terminals (left), and with bath TTX (right) and D. Amplitude of evoked sEPSC from layer 5 PNs (Control, Baseline, Mean ± SD pA, 128.3 ± 132, n = 12; Tinnitus 383 ± 224, n = 14; t (24) = 3.44, *p* = 0.002, *t*-test). ** *p* < 0.01.

### A1 layer 5 PNs show tinnitus-related decreases in GABAergic inhibitory input

Whole-cell patch-clamp electrophysiology from 10 control, 10 ET, and 5 ENT rats was used to examine tinnitus-related changes in the physiology of A1 layer 5B PNs. Increased A1 excitability may, in part, reflect direct and/or indirect loss of synaptic inhibition within A1. SOM, PV and VIP interneurons represent the primary sources of GABAergic inhibition in A1 microcircuitry (Gonchar *et al*., 2008; Blackwell & Geffen, 2017). Spontaneous GABAergic inhibitory currents (sIPSCs) were collected from layer 5 PNs in continuous whole-cell recordings from controls and animals with behavioral evidence of tinnitus (Fig. 4A). Recordings were collected in the presence of bath DNQX and AP5 to block the influence of AMPA and NMDA receptors, respectively. There was a significant tinnitus-related decrease in sIPSCs frequency from layer 5 PNs in ET animals, while layer 5 PNs from ENT animals showed no decrease in sIPSC frequency when compared to control animals (Fig. 4A, B). ET animals showed a significant linear relationship between the decrease in sIPSC frequency and normalized tinnitus score (z-score), which was not seen for control and ENT animals (Fig. 4C). In addition, layer 5 PNs in ET animals showed a significant decrease in sIPSCs amplitude, while no significant difference in sIPSC amplitude was observed in recordings from layer 5PNs from ENT animals. (Fig. 4D). We measured tinnitus-related changes in intrinsic/ambient GABA inhibition in layer 5 PNs by blocking GABA_A_R responses with gabazine. Bath application of gabazine induced a rapid tonic change in holding current in recordings from PNs from control and ENT animals (Fig. 4E, F). This was interpreted as reflecting intrinsic GABAergic tonic inhibitory current mediated via extrasynaptic GABA_A_Rs. PNs from ET animals showed a significant tinnitus-related loss of tonic inhibition revealed by gabazine (Fig. 4E, F). This significant tinnitus-related loss of intrinsic GABA_A_ mediated tonic inhibition at layer 5 PNs likely reflects decreased availability of ambient GABA or loss of extrasynaptic receptors. Miyakawa et al., found a significant tinnitus-related loss of the GABA synthetic enzyme GAD_65_, suggesting a possible decrease in the availability of ambient GABA in animals with evidence of tinnitus (Miyakawa *et al*., 2019). We used fluorescence *in situ* hybridization to identify specific populations of excitatory and inhibitory neurons in A1 from tinnitus and control rats. There were no significant tinnitus-related changes in the density of PV, SOM and VIP neurons or in excitatory (VGLUT1 positive) neurons across A1 layers (Table 2). When layer-specific changes were examined, only layer 4 PV cells from rats with behavioral evidence of tinnitus showed significant tinnitus-related decreases in cell density. Prior findings examining the impact of noise-exposure on PV cell numbers have been inconsistent, with some studies showing no change in PV cell density while other studies showed decreased or increased PV cell density (Nguyen *et al*., 2017; Liu *et al*., 2018; Miyakawa *et al*., 2019; Deng *et al*., 2020).

**Figure 4:**
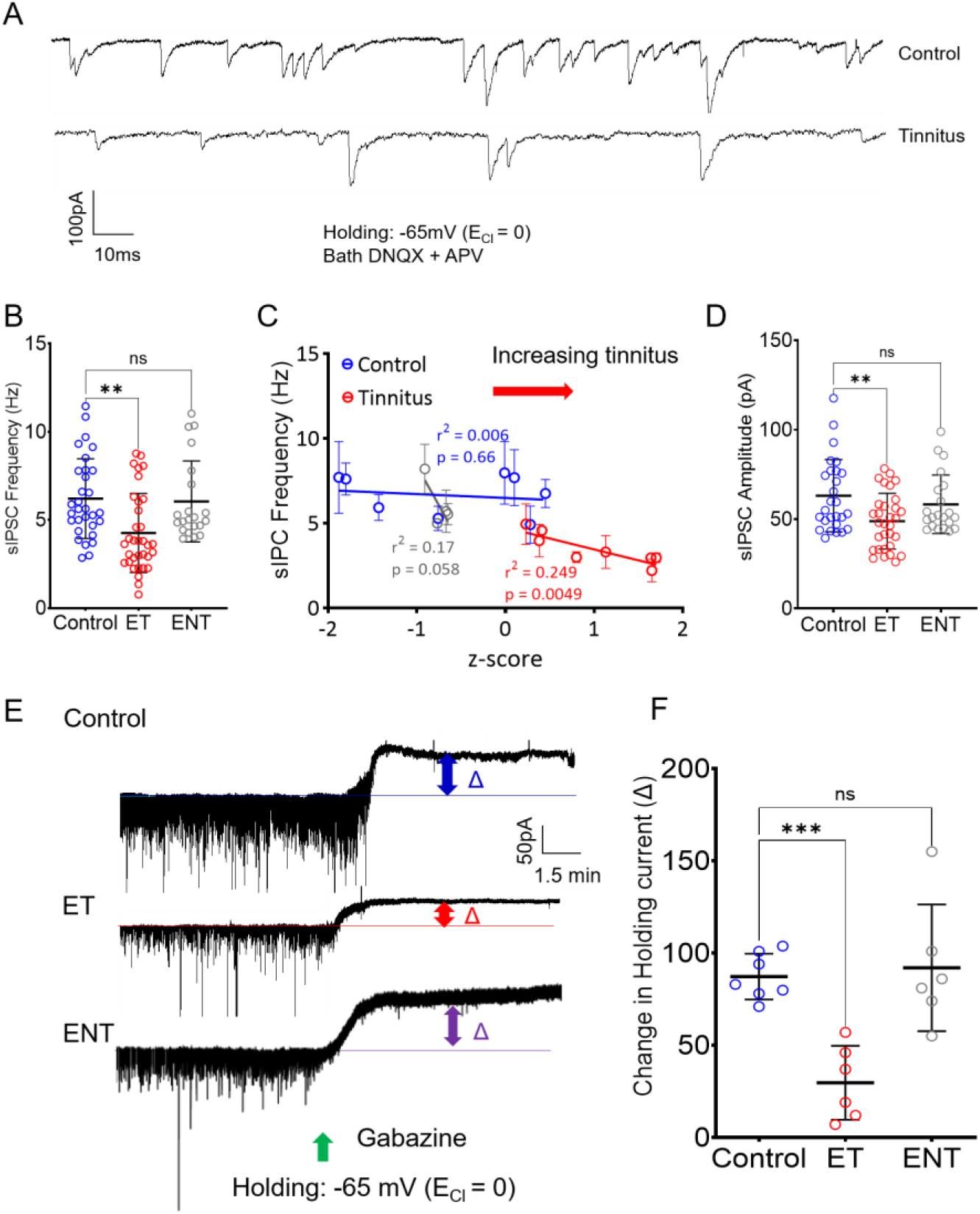
Tinnitus-related loss of GABAergic input onto layer 5 PNs. A. Exemplar traces of sIPSCs recorded from layer 5 PNs from control (top) and tinnitus (bottom) animals. B. Frequency of sIPSCs recorded from layer 5 PNs from control, ET and ENT animals (Control, Mean ± SD Hz, 6.2 ± 2.26, n = 31; ET 4.25 ± 2.24, n = 34; ENT, 6.04 ± 2.29, n = 21 F(2,83) = 7.16, *p* = 0.0013, *p* = 0.0017 control-ET, *p* = 0.9 control *vs ENT*,One-way ANOVA with Bonferroni post-hoc test). C. Plot of sIPSC frequency from layer 5 PNs in control, ET and ENT rats against their normalized tinnitus score (z-score^#^) ((Control, F(1, 30) = 0.18, p = 0.66, r ^2^ = 0.006, ET, F(1, 25) = 8.29, *p* = 0.008, r^2^ = 0.25, ENT, F(1,19) = 4.12, *p* = 0.057, r^2^ = 0.17)). D. Average peak amplitude of sIPSCs from layer 5 PNs from control vs ET animals (Control, Mean ± SD pA, 61.24 ± 20.86, n = 20; ET, 48.73 ± 17.38, n = 20; ENT, 58.24 ± 16.37, n = 21, F(2, 76) = 4.9, *p* = 0.0094, p = 0.0056 control-ET, *p* = 0.69 control-ENT, One-way ANOVA with Bonferroni post-hoc test). E. Exemplar traces showing change in GABA_A_R mediated tonic current in layer 5 PNs from control (top) and tinnitus (bottom) animals. F. Quantification of tonic current (Control, Mean ± SD pA, 87.23 ± 12.43, n = 7; ET, 29.67 ± 20.04, n = 6; ENT, 92.00 ± 34.35, n = 6, F(2, 16) = 13.34, *p* = 0.0004, *p* = 0.0009 control-ET, *p* = 0.9 control-ENT, One-way ANOVA with Bonferroni post-hoc). * *p* < 0.05 **, < 0.01, ***, < 0.001. ^#^ See methods for z-score normalization.

### Tinnitus-related loss of GABA_A_R function

Tinnitus-related decreases in sIPSC amplitude may reflect changes in postsynaptic GABA_A_R composition and/or function. GABAergic currents were examined in recordings from layer 5 PNs in response to puffed GABA (50 μM) 30 μm from the PN being studied (Fig. 5A, B). PNs recorded from animals with behavioral evidence of tinnitus showed significantly smaller GABA evoked IPSCs when compared to PNs recorded from control and ENT animals (Fig. 5C). These findings from layer 5 PNs, suggest tinnitus-related changes in the number and/or subunit composition postsynaptic GABA_A_R. Closer examination of GABA evoked postsynaptic currents showed significantly longer/slower rise-times (Fig. 5D) in PNs from ET animals compared to control and ENT rats. Likewise, changes in GABA evoked decay/fall-times (Fig. 5E) times were significantly different between ET animals compared to control and ENT animals. Altered IPSC rise-fall characteristics have previously been shown to reflect synaptic GABA_A_R subunit changes (Schofield & Huguenard, 2007).

**Figure 5:**
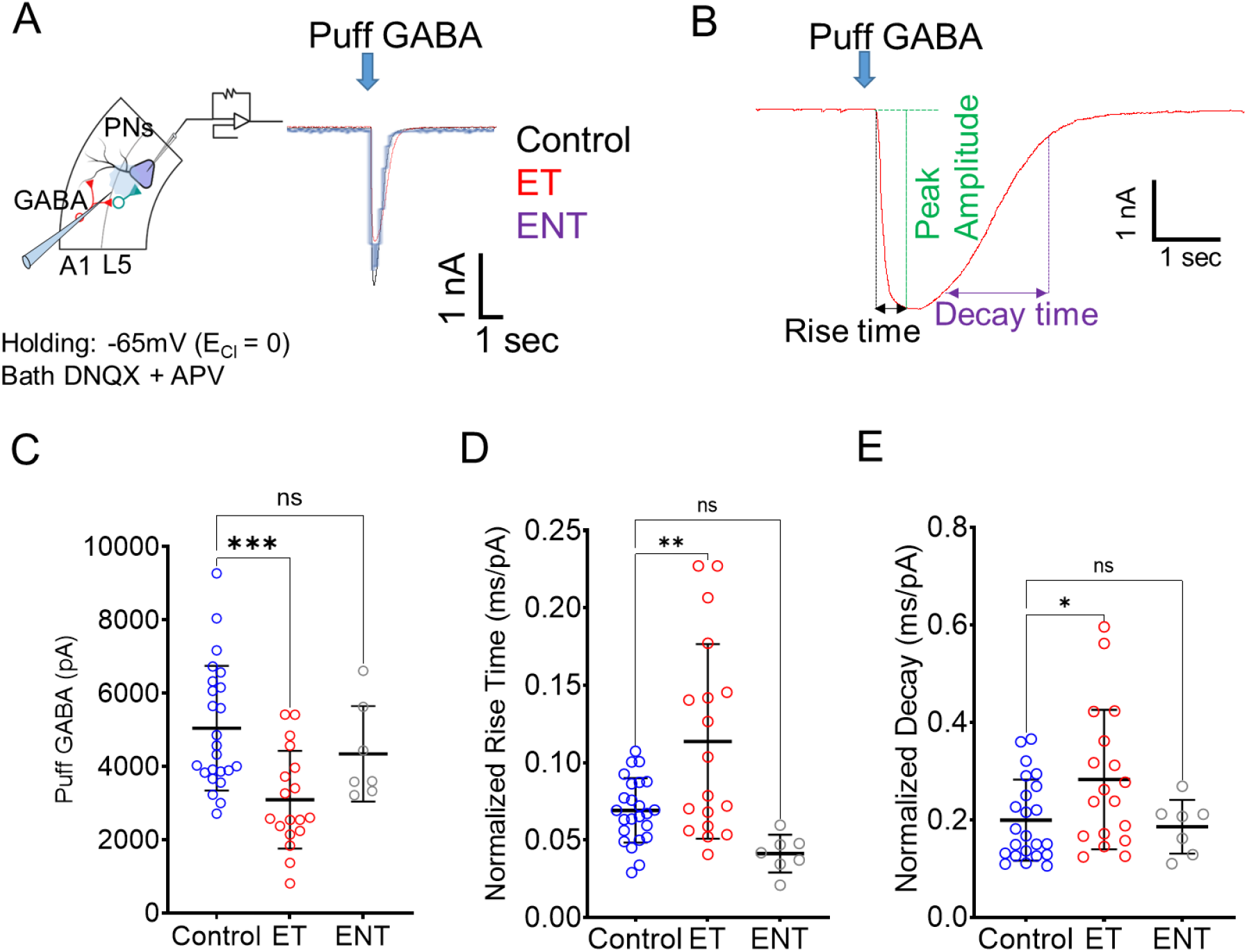
Tinnitus-related disruption in GABA_A_ receptor pharmacokinetics in layer 5 PNs: A. Illustration showing whole-cell patch-clamp recordings from layer 5 PNs while puffing GABA (left); exemplar traces (right) in response to puffed GABA on layer 5 PNs from control (black) and tinnitus (red) animals. B. Measurements of evoked IPSCs in response to puffed GABA, peak amplitude (green), rise time (black) and decay time (purple). C. Peak amplitude of evoked IPSCs to puffed GABA in layer 5 PNs (Control, Mean ± SD pA, 5043 ± 1700, n = 24; ET, 3153 ± 1708, n = 18; ENT, 4343 ± 1305, n = 7, F(2, 46) = 8.44, *p* = 0.0008, *p* = 0.0003 control-ET, *p* = 0.58 control-ENT, Two-way ANOVA with Bonferroni post-hoc test). D. Rise-time normalized to peak amplitude in control and ET animals (Control, Mean ± SD ms/pA, 0.069 ± 0.02, n = 24; ET, 0.11 ± 0.06, n = 18; ENT, 0.04 ± 0.012, n = 7, F(2, 46) = 9.92, *p* = 0.0003, *p* = 0.0023 control-ET, *p* = 0.22 control-ENT, One-way ANOVA with Bonferroni post-hoc test). E. Decay-time normalized to peak amplitude in control and ET animals (Control, Mean ± SD ms/pA, 0.19 ± 0.08, n = 24; ET, 0.28 ± 0.14, n = 18; ENT, 0.18 ± 0.05, n = 7, F(1, 46) = 2.6, *p* = 0.035 control-ET, *p* = 0.9 control-ENT,One-way ANOVA with Bonferroni post-hoc test). * *p* < 0.05 **, <0.01, *** <0.001.

### Extrasynaptic GABA_A_R function remains relatively intact in tinnitus animals

As described above and in Figure 4E, extrasynaptic GABA_A_Rs (α_4_δ) enable neurons to sense ambient extrasynaptyic GABA concentrations reflected by generation of a tonic inhibitory current (Belelli *et al*., 2009). Recordings from A1 layer 5 PNs showed tinnitus-related reductions in the endogenous GABA_A_R inhibitory current mediated by extra-synaptic GABA_A_R blockade (Fig. 6A). We asked if this tinnitus-related change in tonic GABA_A_R current was due to loss of extrasynaptic GABA_A_Rs or a loss of ambient GABA? Bath application of the selective α_4_δ GABA_A_R agonist gaboxadol (GBX, 5 μM, 5 min) was used to evoke tonic GABA_A_R mediated currents that plateaued after few seconds (Fig. 6A). Figure 6B shows nonsignificant tinnitus-related changes in GBX evoked GABA_A_R tonic inhibitory current which suggested that tinnitus-related changes in the numbers of extrasynaptic GABA_A_Rs was unlikely. These results were further supported in a saturation [^3^H]GBX binding study comparing control and noise-exposed animals using the same noise-exposure paradigm as in the present study (Fig. 6C). Consistent with the physiology, binding in A1 contralateral to the exposed ear showed only non-significant changes in the number of [^3^H]GBX binding sites (B_max_) and in the dissociation constant (Kd) (Fig. 6, E). Together, these findings suggest the observed loss of tonic/intrinsic GABA_A_R inhibition in layer 5 PNs from tinnitus rats was likely due to lowered availability of ambient GABA in A1 as suggested by previous human and animal studies (Llano *et al*., 2012; Sedley *et al*., 2015; Deng *et al*., 2020).

**Figure 6:**
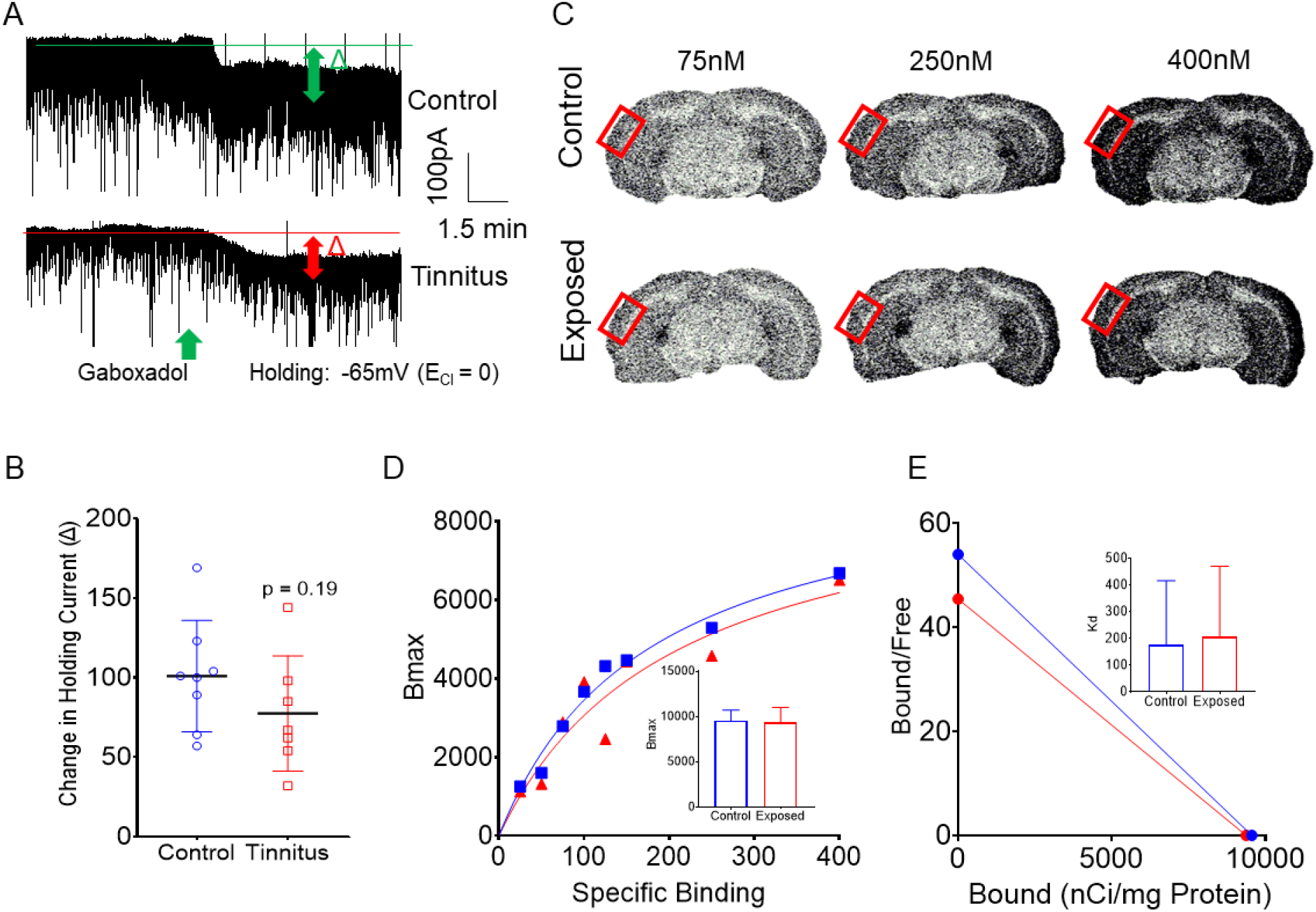
A. Exemplar whole-cell recordings showing extrasynaptic GABA_A_R mediated tonic current in layer 5 PNs of control (top) and tinnitus (bottom) animals following bath application of gaboxadol (GBX). B. Tinnitus-related change in holding current in response to bath GBX (Control, Mean ± SD pA, 100.9 ± 34.98, n = 8 neurons, 2 animals; Tinnitus, 77.43 ± 34.11, n = 7 neurons, 2 animals; t(13) = 1.27, *p* = 0.225, *t*-test). C. Exemplar autoradiographic images showing [^3^H]GBX binding in coronal brain sections containing A1 (red box) from control (top) and similarly noise-exposed (bottom) animals at 75, 250 and 400 nM concentration. D. Saturation binding curves show maximum binding sites in the A1 in control vs noise-exposed animals (Control, Mean ± SD nCi/mg protein, 9561 ± 1166, n = 30 samples, 10 animals; Tinnitus, 9376 ± 1627, n = 15 samples, 5 animals; t(43) = 0.45, *p* = 0.66, *t*-test). E. Calculation of dissociation constant (*Kd*) as a function of binding affinity show no noise-exposure related changes (Control, Mean ± SD nM 177.1 ± 240.1, n = 30 samples, 10 animals; Tinnitus, 206.4 ± 264.8, n = 15 samples, 5 animals; t (43) = 0.37, *p* = 0.71, *t*-test).

### A1 VIP neurons show tinnitus-related increased excitability

Cortical VIP neurons play a key role in regulating the activity of PNs. The canonical inhibitory microcircuitry of A1 describes inhibitory VIP neurons as projecting onto inhibitory PV and SOM neurons, which in turn provide for disinhibitory regulation of layer 5 PNs (Mesik *et al*., 2015; Askew *et al*., 2019). We posited that tinnitus-related increases in VIP neuronal activity would inhibit PV and SOM neurons resulting in disinhibition of layer 5 PNs. To test this hypothesis, VIP-Cre LE rats were bred with a TdTomato reporter line of LE rats to obtain offspring expressing TdTomato in VIP-positive neurons (Fig. 7A). Expression of tdTomato VIP neurons was immunohistochemically confirmed using an anti-VIP antibody (Fig. 7B, C). VIP x tdTomato rats were inserted into the previously described, noise-exposure/training/tinnitus testing protocol and classified as non-exposed controls or animals with behavioral evidence of tinnitus (ET). VIP neurons were identified in A1 slices using fluorescence microscopy and studied using whole-cell patch-clamp recordings as above. The intrinsic properties and excitability profile of VIP neurons from control animals were compared with neurons from ET rats. VIP neurons from ET animals were significantly depolarized compared to controls (Fig. 7D, E). VIP neurons from ET animals were highly excitable at significantly lower injected currents/RC (current required to generate action potential) when compared to VIP neurons from control animals (Fig. 7E). Injection of twice the threshold current (2XRC) evoked significantly greater numbers of action potentials (APs) from VIP neurons in ET animals (Fig. 7G).

**Figure 7:**
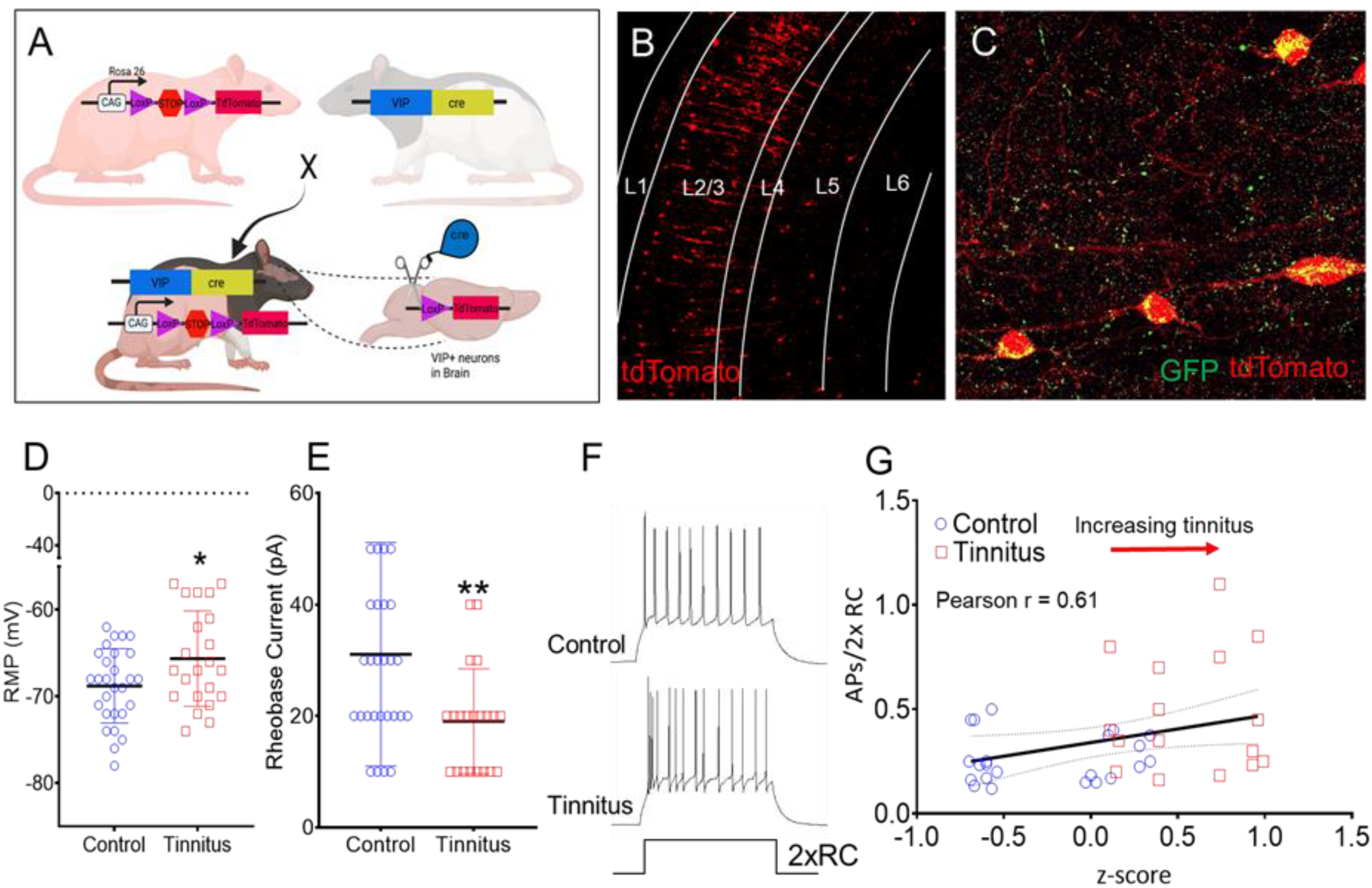
VIP neurons show tinnitus-related increased excitability. A. Illustration showing the generation of *VIP^Cre^:Rosa26^tdTomato^* rats by breeding VIP-Cre rats with tdTomato reporter rats. B, C. *VIP^Cre^:Rosa26^tdTomato^* rats express fluorescent tdTomato protein in VIP+ rats which was validated by using anti-VIP antibody in A1 sections. D. Tinnitus-related change in VIP resting membrane potential (Control, Mean ± SD mV, −68.29 ± 5.03, n = 24; Tinnitus, −63.86 ± 6.2, n = 14; t(36) = 2.4, *p* = 0.021, *t*-test) and E. Tinnitus-related change in VIP rheobase current (Control, mean ± SD pA, 28.33 ± 21.4, n = 24; Tinnitus, 16.92 ± 7.5, n = 13; t(31) = 2.17, *p* = 0.037, *t*-test) recorded from A1 VIP+ neurons in control and tinnitus rats. F. Exemplar traces showing the VIP neurons firing/APs upon injection of twice rheobase current (2 x RC). G. Plot of the number of action potentials (APs) generated by injection of twice the rheobase current in VIP neurons from all control and tinnitus rats plotted against their normalized tinnitus score (z-score^#^) (Control, Mean ± SD APs/2xRC, 0.259 ± 0.05, n = 20; Tinnitus, 0.479 ± 0.15, n = 17; Pearson r = 0.61, *p* = 0.033). * *p* < 0.05. ^#^ See methods for z-score normalization.

## Discussion

The present studies found significant tinnitus-related maladaptive plastic changes in A1 layer 5 PNs and VIP inhibitory neurons in an established behavioral rat model of tinnitus. 1) Layer 5 PNs from ET animals were significantly depolarized at rest, reflecting a disrupted excitatory-inhibitory balance. 2) Layer 5 PNs displayed increased frequency of sEPSCs and decreased frequency of inhibitory currents (sIPSCs) with these tinnitus-related changes directly correlated to the severity of the rat’s tinnitus behavioral score. 3) Postsynaptic GABA_A_Rs showed tinnitus-related reductions in their responses to GABA, suggestive of decreases in GABA_A_R expression or altered subunit make-up of GABA_A_Rs (Wafford *et al*., 1993). 4) Layer 5 PNs showed enhanced thalamocortically evoked polysynaptic EPSCs in ET animals. Previous studies describe tinnitus-related changes suggestive of increases in the number of active ascending MGB channels, increased bursting indicative of thalamocortical dysrhythmia or tinnitus-related increases in neurotransmitter availability/release onto postsynaptic sites (Kalappa *et al*., 2014; De Ridder *et al*., 2015; Elgoyhen *et al*., 2015; Caspary & Llano, 2017). 5) Columnar VIP neurons, thought to modulate PN excitability via disinhibition, showed tinnitus-related increases in excitability including reduction in resting membrane potential (depolarized state) and 6) tinnitus-related increases in rheobase current-evoked action potentials. Numbers of evoked action-potentials from VIP neurons were directly correlated with the severity of the tinnitus behavioral score. We hypothesize that tinnitus-related increases in VIP excitability results in inhibition of SOM and PV inhibitory interneurons, disinhibiting layer 5 PNs. Therefore, A1 VIP neurons may form an important link in our understanding of tinnitus-related maladaptive plastic changes within A1. Understanding the impact of tinnitus on the synaptic and intrinsic properties of A1 VIP neurons is important to better appreciate circuit changes underpinning tinnitus pathology. These same PN and VIP changes observed here have been previously detailed in a mouse model of neuropathic pain in the somatosensory cortex (Cichon *et al*., 2017). Collectively these finding suggest that tinnitus and chronic pain may share common cortical pathology.

The present findings are, in part, consistent with the maladaptive plasticity hypothesis of Turrigiano and colleagues, which details that activity deprivation induces homeostatic changes in target neurons (Turrigiano *et al*., 1998). Noise-exposure used to induce tinnitus in the present study resulted in a permanent threshold shift at frequencies adjacent to the noise-exposure frequency. Accordingly, long-term sensory damage/deprivation is likely to initiate compensatory homeostatic mechanisms generally seen as increased spontaneous and driven activity, bursting and cross-connectivity in cochlear nucleus, inferior colliculus, auditory thalamus and auditory cortex (Brozoski *et al*., 2002; Bauer *et al*., 2008; Roberts *et al*., 2010; Noreña & Farley, 2013; Sedley, 2019; Shore & Wu, 2019; Henton & Tzounopoulos, 2021; Zhai *et al*., 2021; McGill *et al*., 2022).

In A1, tinnitus-related increases in sEPSC frequency and decreases in sIPSC frequency impacted layer 5 PN excitability changes and were directly correlated with tinnitus severity in behaviorally measured tinnitus (Figs. 2, 4), while layer 5 PNs from ENT animals showed no change in excitatory input. Tinnitus-related changes in excitatory/inhibitory homeostasis are consistent with previously described tinnitus-related plastic changes in central auditory neurotransmission (Noreña & Farley, 2013; Auerbach *et al*., 2014). Layer 5 PNs provide excitatory feedback to the MGB and extra-lemniscal IC as well as local cortical circuits (Williamson & Polley, 2019; Kommajosyula *et al*., 2021; Lesicko & Geffen, 2022). Tinnitus-related increases in layer 5 excitability theoretically could increase neuronal excitation at these target structures, potentially initiating reciprocal extra-lemniscal feed-forward excitation to A1 (Syka & Popelář, 1984; Stebbings *et al*., 2014; Williamson & Polley, 2019; Blackwell *et al*., 2020).

Miyakawa and colleagues found tinnitus-related decreases in A1 levels of the GABA synthesizing enzyme, GAD_65_ (Miyakawa *et al*., 2019). Consistent with the present findings, Yang and colleagues (2011) working in a mouse model of tinnitus, found that A1 layer 2/3 PNs showed tinnitus-related down-regulation of GABA synthesis accompanied by increases in sEPSCs and decrease in sIPSCs (Yang *et al*., 2011). In mouse, noise exposure-related losses of PV interneurons, responsible for feed-forward inhibition, were hypothesized to decrease GABA release onto PNs in A1 (Miyakawa *et al*., 2019). Here in the rat model, we observed a significant loss of PV neurons in layer 4 and no significant loss of PV neurons across other layers of A1 (Table 2) but found a significant tinnitus-related down-regulation of sIPSCs onto layer 5 PNs. A significant, and likely compensatory, increase in sEPSC frequency onto layer 5 PNs also correlated to the behavioral measure of tinnitus across groups (Fig. 2E). Changes in sIPSC or sEPSC frequency are believed to reflect changes in input neurons activity. While amplitude changes are generally considered to reflect both postsynaptic changes and changes in neurotransmitter release from the input neurons (Bruns & Jahn, 1995). Tinnitus-related increases in sEPSC frequency suggested that the layer 5 PNs are receiving a greater level of excitatory input from excitatory connections within cortex: cortico-cortical, A1 columnar or non-A1 (Kalappa *et al*., 2014; De Ridder *et al*., 2015; Elgoyhen *et al*., 2015). Optogenetic stimulation of thalamocortical terminals evoked significantly larger EPSCs from ET animals, which could reflect increased glutamate vesicular release, upregulation of AMPA/NMDA receptors or simply reflect tinnitus-related losses of PN inhibitory input (Murthy *et al*., 2001; Yang *et al*., 2011; De Ridder *et al*., 2015). A significantly larger portion of the thalamocortical EPSCs was mediated via polysynaptic mechanisms, in part, supporting tinnitus-related increases in excitability throughout A1 layers (Fig. 3C) (Slater *et al*., 2019). The axonal potassium channel blocker 4-aminopyridine, usually used to increase terminal release excitability in experiments involving TTX (Shu *et al*., 2007; Cho *et al*., 2013), was deemed not necessary in our studies, given a strong LED-induced postsynaptic currents were evoked, likely due to the high release probability at thalamocortical terminals (Gil *et al*., 1999; Slater *et al*., 2019).

A1 layer 5 microcircuitry involves several types of excitatory pyramidal neurons and at least 4 different types of inhibitory interneurons. Layer 5 PNs receive local inhibitory input from nearby interneurons, primarily from PV and SOM interneurons, known to regulate their excitability (Letzkus *et al*., 2011; Mesik *et al*., 2015; Askew *et al*., 2019). Findings in present studies show tinnitus-related losses of pre- and postsynaptic GABAergic inhibition onto layer 5 PNs, consistent with the studies in layer 2/3 of A1 by Miyakawa and colleagues (Miyakawa *et al*., 2019). Similar to findings in a neuropathic pain model, A1 layer 5 neurons were found to receive significantly lower levels of inhibition as a result of tinnitus-related plastic changes of normal PV and SOM function (Fig. 4E) (Cichon *et al*., 2017). VIP neurons from ET animals showed a significant increase in the numbers of action potentials evoked by depolarizing electrical stimulation which directly correlated to the animal’s tinnitus score. This VIP hyperexcitability was in part due to the tinnitus-related depolarized RMP likely again reflecting decreased endogenous levels of GABA (Fig. 7G). Activation of VIP neurons is known to inhibit the activity of PV and SOM inhibitory interneurons, resulting in decreased GABA_A_R mediated tonic and phasic inhibition at the level of layer 5 PNs (Letzkus *et al*., 2011; Mesik *et al*., 2015; Cichon *et al*., 2017; Askew *et al*., 2019). Tinnitus-related increased excitability of VIP neurons would underpin the observed decreased feedforward inhibition via PV and SOM neurons, likely depolarizing layer 5 PNs (Fig. 2B) and similar to what was seen in somatosensory cortex by Cichon et al. (2017).

We found that PNs from ET animals were less sensitive to puff-applied GABA producing smaller amplitude/peak IPSCs and suggesting a significant loss of postsynaptic GABA_A_R function. Previously described tinnitus-related loss of GAD_65_ suggests decreased GABA synthesis, which has been shown to alter GABA_A_R number and/or GABA_A_R pharmacology via subunit switching (Wafford *et al*., 1993; Murthy *et al*., 2001). Extrasynaptic α_4_β_2_δ-containing GABA_A_Rs are present in the neocortex and mediate robust non-desensitizing tonic inhibition (Drasbek & Jensen, 2005; Caspary & Llano, 2017). The observed tinnitus-related loss of tonic GABA_A_R current was likely due to a loss of ambient GABA, since the present patch-clamp and receptor binding studies found no significant tinnitus-related changes suggestive of a loss in extrasynaptic GABA_A_R number (Fig. 6C-E).

Tinnitus not only affects the auditory pathways, it also affects various attentional and limbic/emotional systems (Kraus & Canlon, 2012; Chen *et al*., 2017; Besteher *et al*., 2019). A1 layer 5 PNs recorded in the present study may make multiple excitatory connections to subcortical centers such as IC, MGB, amygdala or intra-cortical targets (Asokan *et al*., 2018; Baker *et al*., 2018). Increased activity of layer 5 PNs may worsen tinnitus pathology by various mechanisms including facilitation of: 1) the activity of trauma-associated limbic regions; 2) the pathological reverberant feedback loop or pathological synchrony between IC, MGB and A1; and 3) amplifying cortical responses to sub-perceptual inputs, and/or combinations of all these possibilities.

In conclusion, in an established tinnitus model, the present study describes tinnitus-related pre- and postsynaptic changes in the physiology of A1 layer 5 pyramidal and VIP neurons from ET animals but not from ENT rat A1. Tinnitus-related increases in excitability of layer 5 PNs was accompanied by increased excitability of VIP interneurons. Future studies will examine tinnitus-related changes in PV and SOM inhibitory interneurons while considering pharmacologic agents targeting VIP excitability.

## Data availability statement

All data supporting the results have been included in the manuscript and figures. Additional data in support of the findings of this study are available from the corresponding author, upon request.

## Acknowledgement

This work was supported by Department of Defense Award W81XWH1910017 to DMC, NIH DC015388 to TAH, and NIH DC016599 and DC013073 to DAL. We thank Dr. Thomas Brozoski, Dr. Brandon Cox, and Kurt Wisner for assistance with the animal model and Dr. Benjamin Richardson for sharing his knowledge of electrophysiological data analysis.

## Author Contributions

Electrophysiology and associated imaging studies carried out in DMC Neurobiology laboratories at SIU-SM. FISH studies by TAH at Vanderbilt University; MG: study design, data acquisition, data analysis, data interpretation, manuscript drafting and editing; RC: data analysis, interpretation manuscript editing; LL: Receptor binding, IHC, data collection and interpretation, manuscript editing; KB: Operant behavior, generation, training and testing tinnitus subjects; TAH: generation, interpretation analysis of FISH data, manuscript editing; DAL: study design, data interpretation, manuscript editing; DMC: study design, data analysis, data interpretation, manuscript editing. All authors have approved final version of the manuscript. DMC agrees to be accountable for all aspects of the work and will ensure that questions related to the accuracy or integrity of any part of the work are appropriately investigated and resolved.

